# Genotypic frequencies at equilibrium for polysomic inheritance under double-reduction

**DOI:** 10.1101/532861

**Authors:** Kang Huang, Tongcheng Wang, Derek W. Dunn, Pei Zhang, Xiaoxiao Cao, Rucong Liu, Baoguo Li

**Affiliations:** Shaanxi Key Laboratory for Animal Conservation, College of Life Sciences, Northwest University, Xian, China, 710069.; Center for Excellence in Animal Evolution and Genetics, Chinese Academy of Sciences, Kunming, China 650223.

**Keywords:** Polysomic inheritance, double-reduction, genotypic frequency, inbreeding coefficient, heterozygosity

## Abstract

Polyploids are organisms whose genomes consist of more than two complete sets of chromosomes. Both autopolyploids and allopolyploids may display polysomic inheritance. A peculiarity of polysomic inheritance is multivalent formation during meiosis resulting in double-reduction, which occurs when sister chromatid fragments are segregated into the same gamete. Double-reduction can result in gametes carrying identical-by-descent alleles and slightly increasing homozygosity. This will cause the genotypic frequencies to deviate from expected values and will thus bias the results of standard population genetic analytical methods used in molecular ecology and selective breeding. In this study, we extend existing double-reduction models to account for any even level of ploidy, and derive the symbolic expressions for genotypic frequencies via two methods. Inbreeding coefficients and heterozygosity under double-reduction and inbreeding are also calculated. Numerical solutions obtained by computer simulations are compared with analytical solutions predicted by the model to validate the model.

## INTRODUCTION

Polyploids are organisms whose genomes consist of more than two complete sets of chromosomes (Madlung, 2013). They represent a significant proportion of plant species, with 30-80% of angiosperm species showing polyploidy (Burow *et al.*, 2001) and most lineages showing evidence of paleoploidy (Otto, 2007). Polyploid plants can arise spontaneously in nature by several mechanisms, including meiotic or mitotic failures, and fusion of unreduced gametes (Comai, 2005).

There are two distinct mechanisms of genome duplication that result in polyploidy: allopolyploidy and autopolyploidy. Autopolyploids are usually thought to arise within a species by the doubling of structurally similar homologous genomes, whereas allopolyploids arise via interspecific hybridization and subsequent doubling of non-homologous genomes (Parisod *et al.*, 2010). Both autopolyploids and allopolyploids can be found among both wild and domesticated plant species. Although rare, polyploidy is also found in a few species of vertebrates such as some salmonid fish (Limborg *et al.*, 2017), the weather loach (*Misgurnus anguillicaudatus*) (Zhou *et al.*, 2016), the common carp (*Cyprinus carpio*) (David *et al.*, 2003), and the African clawed frog (*Xenopus laevis*) (Session *et al.*, 2016).

In autopolyploids, more than two homologous chromosomes can pair at meiosis, resulting in the formation of multivalents and polysomic inheritance (Rieger *et al.*, 1968). A peculiarity of polysomic inheritance is the possibility that a gamete inherits a single gene copy twice, termed double-reduction (Butruille and Boiteux, 2000). For example, an autotetraploid individual *ABCD* produces a gamete *AA*. In prophase I, crossovers can happen between the locus and the centromere, resulting in an exchange of chromatid fragments between pairing chromosomes. In a multivalent configuration, the separation of chromosomes can be either disjunctional or nondisjunctional (de Silva *et al.*, 2005). For nondisjunctional separation, chromosomes that were previously paired in Prophase I are segregated into a same secondary oocyte. Double-reduction occurs when fragments of sister chromatids are further segregated into a single gamete.

Double-reduction arises from a combination of three major events during meiosis: crossing-over between non-sister chromatids, an appropriated pattern of disjunction, and the subsequent migration of the chromosomal segments carrying a pair of sister chromatids to the same gamete (Darlington, 1929; Haldane, 1930). Multivalent chromosome pairing during meiosis can also occur in allopolyploids, resulting in a mixed inheritance pattern across loci in the genome, termed segmental allopolyploidy (Stebbins, 1950).

Geneticists have developed several mathematical models to simulate the double-reduction. For instance, for tetrasomic inheritance, the rate of double-reduction *α* is assumed to have a minimum value of 0 under *random chromosome segregation* (RCS, Muller, 1914). This increases to 1/7 with *pure random chromatid segregation* (PRCS, Haldane, 1930), and to 1/6 with *complete equational segregation* (CES, Mather, 1935). A diagram of these models can be found in Figure 1.

**Figure 1.**
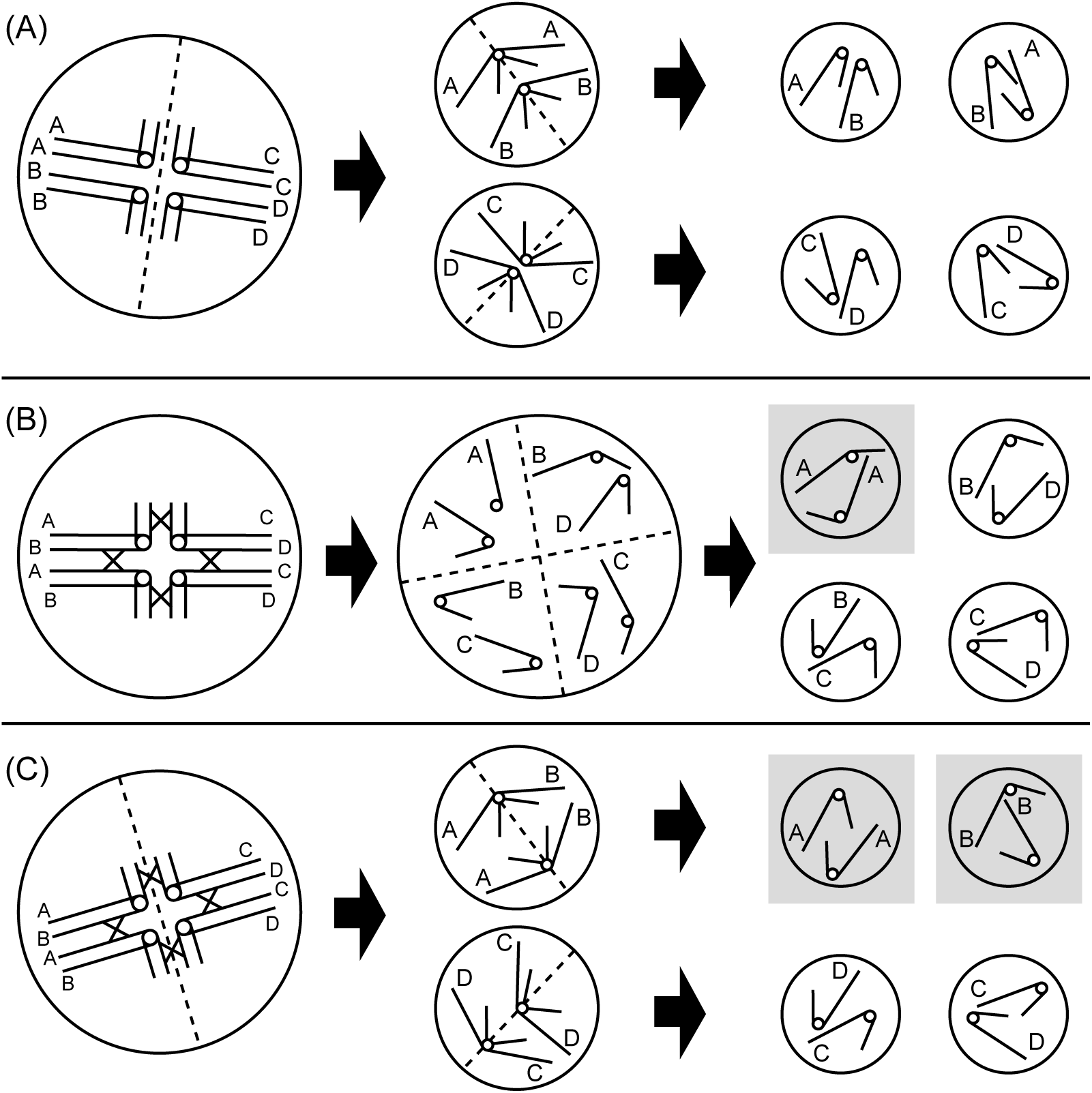
Diagrams of double-reduction models under tetrasomic inheritance. The leftmost column shows the primary oocytes, the middle column shows the secondary oocytes (A and C) or tetrad (B) and the rightmost column shows the gametes. The gametes with a gray background carry identical-by-double-reduction alleles. Dashed lines denote cellular fission, solid lines denote the arms of the chromosome, and the circles connecting solid lines denote the centromere. The target locus is located in the long arm of the chromosome and the identical-by-descent alleles are denoted by the same letter. (A) Random chromosome segregation (RCS) ignores the crossover between the target locus and the centromere (Muller, 1914). In the absence of crossing over, gametes may originate from any combination of homologous chromosomes, and two sister chromatids never sort into the same gamete (Parisod *et al.*, 2010); (B) Pure random chromatid segregation (PRCS) accounts for the crossing over between the target locus and the centromere, and assumes the chromatids behave independently and randomly segregate into gametes (Haldane, 1930). When sister chromatids are segregated into the same gamete, double-reduction occurs. The probability that the two chromatids in a gamete are sister chromatids is 4 (the number of sister chromatids pairs) divided by 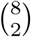 (the number of ways to sample two chromatids from eight chromatids), which is equal to 1/7. (C) Complete equational segregation (CES), homologous chromosomes pair and chromatids exchange via recombination (Mather, 1935). The whole arms of sister chromatids are exchanged into different chromosomes. The probability that two homologous chromosomes within a single secondary oocyte were previously paired at a target locus in Prophase I is 1/3. In this case, these sister chromatid fragments will become segregated in a single gamete at a ratio of 1/2, so the rate of double-reduction is thus 1/6 for tetrasomic inheritance.

Double-reduction results in gametes carrying *identical-by-descent* (IBD) alleles and slightly increases homozygosity (Hardy, 2016). Experiments aimed at estimating the frequency of double-reduction in autotetraploids have yielded values ranging from 0 to almost 0.30 (Wu, Rong Ling and Gallo-Meagher, Maria and Littell, Ramon C. and Zeng, Zhao Bang, 2001). It has a similar effect as inbreeding, and can cause a deviation in genotypic frequencies from Hardy-Weinberg equilibrium (Luo *et al.*, 2006). Fisher (1943) first studied the Mid gene of *Lythrum salicaria*, and produced an analytical method to estimate genotypic frequencies for tetrasomic inheritance for a biallelic locus. Geiringer (1949) extended this work to a multiallelic locus for tetrasomic inheritance. However, the genotypic frequencies for polysomic inheritance for higher levels of ploidy and for a multiallelic locus have been neglected.

Genotypic frequencies are important in population genetics, molecular ecology and molecular breeding, with the *Hardy-Weinberg equilibrium* (HWE) being the basal standard for analytical methods in disomic inheritance. Several methods assume that loci conform to the HWE for the calculations of certain model parameters or statistics, for example, the likelihood of a genotype (population assignment, Bayesian clustering, parentage analysis), the statistics measuring the deviation from expectations (linkage disequilibrium test, differentiation test, heterozygote deficiency/excess test, bottle-neck effect test), and other parameters (e.g. individual inbreeding coefficient, relatedness coefficient, kinship coefficient, QTL mapping). Although some analytical methods have been developed for organisms with polysomic inheritance (e.g. Clark and Jasieniuk, 2011; Hardy and Vekemans, 2002; Huang *et al.*, 2015; Meirmans and van Tienderen, 2004; Pritchard *et al.*, 2000), none of these methods account for double-reduction, and thus in the presence of double-reduction will produce misleading results. For example, in tetrasomic inheritance, *ABCD* × *EFGH* may produce an offspring *AAEE*, and both parents are excluded as the true parents in parentage analysis.

Here, we extend an existing double-reduction model to incorporate the recombination rate into CES, and show two methods to derive the symbolic expressions for genotypic frequencies at equilibrium. To enable other researchers make similar calculations, we make available a C++ source code that enables the calculation of genotypic/phenotypic frequencies of gametes/zygotes at equilibrium to a maximum ploidy level of ten.

## DOUBLE-REDUCTION MODELS

In this paper, we assume that the population size is sufficiently large and there are no differences in selection between both zygotes and gametes. All zygotes will thus have an equal opportunity to mature and reproduce, and all gametes will have an equal opportunity to become fertilized. Therefore, the zygote frequencies are equal to the genotypic frequencies of reproducing individuals, and as a result, we use the zygote frequencies to indicate the genotypic frequencies of reproducing individuals. The double-reduction rates (i.e. alpha) are assumed to be equal among reproducing individuals (e.g. in females and males). Mutation and migration are not included in our model, whereas inbreeding is included for analyzing the influence of inbreeding and double-reduction on the inbreeding coefficient and for estimating heterozygosity. Organisms with odd levels of ploidy are not considered because of their inherent inability to produce euploid gametes — these organisms are sterile.

### Transitional probability from zygotes to gametes

The transitional probability from a zygote to a gamete is the probability that a zygote produces a gamete with a certain genotype, which can be used to find the double-reduction rates of PRCS and to derive the analytical expression of gamete frequencies in the first step of the non-linear method (see Non-linear method section). Such a probability can also be applied to some population genetics studies such as parentage analysis, and the calculation of segregation ratios in mapping populations (see Discussion). If the rate of double-reduction is greater than zero, the gametes will carry *identical-by-double-reduction* (IBDR) alleles. For tetrasomic and hexasomic inheritance, there are only two and three allele copies within a gamete, respectively. Hence, it can be found at most one pair of IBDR alleles within a gamete, and a single parameter can be used to measure the rate of double-reduction.

In polysomic inheritance with a higher level of ploidy, there may be more than one pair of IBDR alleles in a gamete. Therefore it is necessary to add some additional parameters in the segregation ratios. The transitional probability from zygotes to gametes for tetrasomic and hexasomic inheritances at a biallelic locus (the Mid gene in *Lythrum salicaria*) has been studied previously by Fisher and Mather (1943). Here, we extend the work of Fisher and Mather and use a general mathematical expression of this transitional probability. There are at most ⌊*v*/4⌋ pairs of IBDR alleles within a gamete, where *v* is the ploidy level of the resulting zygote. Let *G* and *g* be the genotypes of a zygote and a gamete, respectively, and let *α*_*i*_ be the probability that the gamete carries *i* pairs of IBDR alleles, then 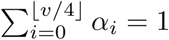 Suppose that *h* is the number of alleles at target locus, then the probability that *G* produces *g* is the following weighted sum:

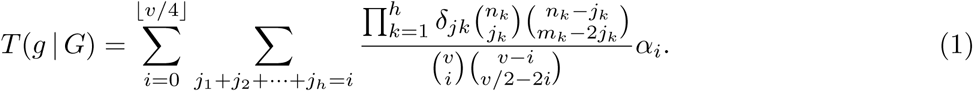

Where *n*_*k*_ and *m*_*k*_ are the numbers of copies of the *k*^th^ allele (say *A*_*k*_) in *G* and *g*, respectively, and *j*_*k*_ is the number of IBDR allele pairs *A*_*k*_-*A*_*k*_ in *g, k* = 1, 2, …, *h*, then 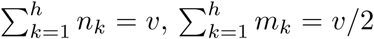, and 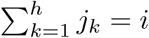.

The symbol *δ*_*jk*_ in Equation (1) is a binary variable, which is used to limit the range of *j*_*k*_, so as to avoid some invalid situations. Note that the combination 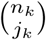 implies 0 ⩽ *j*_*k*_ ⩽ *n*_*k*_, and 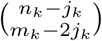 implies 0 ⩽ *m*_*k*_ − 2*j*_*k*_ ⩽ *n*_*k*_ − *j*_*k*_, or equivalently it implies *j*_*k*_ ⩽ *m*_*k*_/2 and *m*_*k*_ − *n*_*k*_ ⩽ *j*_*k*_. Hence *δ*_*jk*_ = 1 if max(0, *m*_*k*_ − *n*_*k*_) ⩽ *j*_*k*_ ⩽ min(*n*_*k*_, *m*_*k*_/2), otherwise *δ*_*jk*_ = 0.

In Equation (1), for every *i* (*i* = 0, 1, 2, …, ⌊*v*/4⌋), two events need consideration. First, is the event that *g* carries *i* pairs of IBDR alleles. Second, is the event that those *i* pairs of IBDR alleles are distributed into *h* kinds of alleles. Because both types of event are mutually exclusive, the products of the probabilities of the two events stated above are summed to obtain *T* (*g* |*G*).

Each chromosome is duplicated during meiosis, so each allele in *G* becomes a pair of duplicated alleles. For the *k*^th^ kind of allele, firstly we sample *j*_*k*_ pairs from the *n*_*k*_ pairs of duplicated alleles *A*_*k*_-*A*_*k*_, then there are 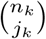 ways (where 0 ⩽ *j*_*k*_ ⩽ *n*_*k*_), and so the first combination in the numerator of Equation (1) is obtained. We then sample the non-IBDR alleles to form *g*. The remaining is *n*_*k*_ − *j*_*k*_ pairs of duplicated alleles *A*_*k*_-*A*_*k*_, which still requires *m*_*k*_ − 2*j*_*k*_ copies of *A*_*k*_. Then we sample *m*_*k*_ − 2*j*_*k*_ pairs of duplicated alleles *A*_k_ -*A*_*k*_, and so there are 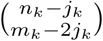 ways (where 0 ⩽ *m*_*k*_ − 2*j*_*k*_ ⩽ *n*_*k*_ − *j*_*k*_), which obtains the second combination in the numerator of Equation (1). Moreover, if we sample one copy of *A*_*k*_ in each pair of the *m*_*k*_ − 2*j*_*k*_ pairs of duplicated alleles, then there are altogether 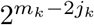 possible combinations. Because each sampling combination is independent for different kinds of alleles, the total number of combinations is the product 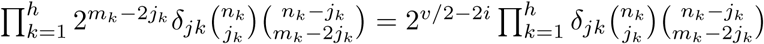

In the following example, we consider the total number of combinations to produce a gamete consisting of *i* pairs of IBDR alleles without accounting for the specific genotypes *G* and *g*. Similar to the above process, we firstly sample *i* pairs from the *v* pairs of duplicated alleles, and these alleles will become the IBD alleles in *g*. We then sample *v*/2 − 2*i* pairs from the remaining *v* − *i* pairs, and then one allele in each pair of the *v*/2 − 2*i* pairs is further sampled to form *g*. The total number of allele combinations is 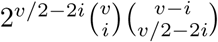 Here the coefficient 2^*v*/2^−^2*i*^ is eliminated in the fraction in Equation (1).

Equation (1) is a general expression of the transitional probability, and can be applied at multiallelic loci and at any even level of ploidy. As an example, the transitional probability from zygotes to gametes in octosomic inheritance at a biallelic locus is shown in Table 1. Another example is shown in Appendix A, which is the derivation of *T* (*BBBB* |*ABBBBBBB*) = (4*α*_0_ + 5*α*_1_ + 6*α*_2_)/8.

**Table 1:**
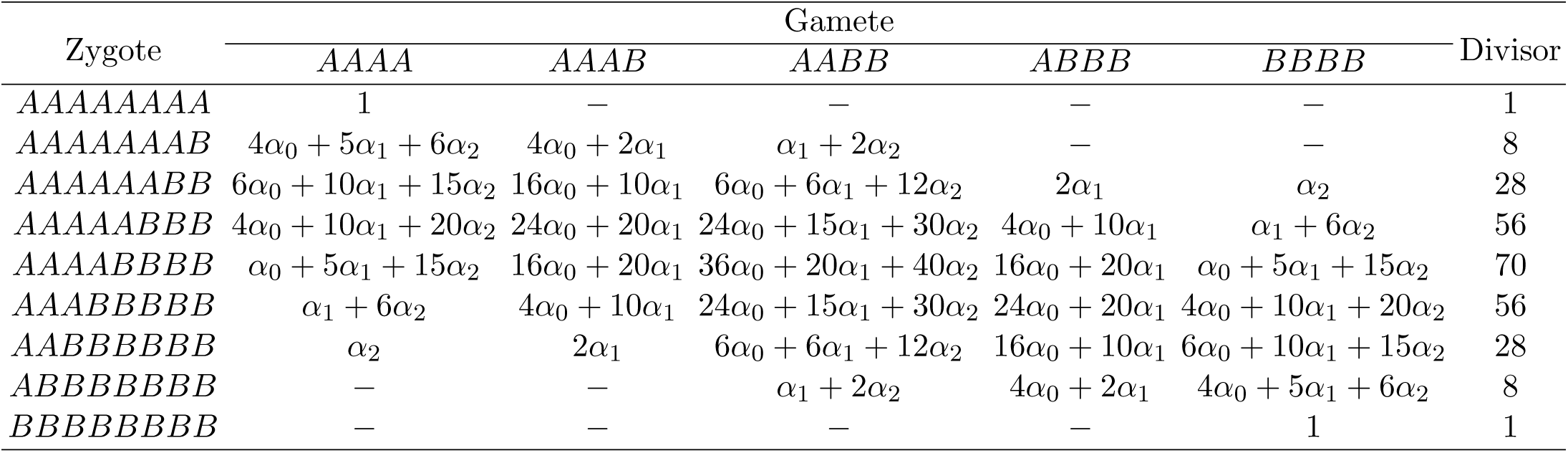
Transitional probability from zygotes to gametes in octosomic inheritance at a biallelic locus

### Random chromosome segregation

The random chromosome segregation model ignores the crossover between the target locus and the centromere, and the rate of double-reduction (*α*) is equal to zero (Muller, 1914). In the absence of crossing over, gametes may originate from any combination of homologous chromosomes, and two sister chromatids never sort into the same gamete (Parisod *et al.*, 2010, Figure 1A). Therefore, genotypic frequencies concur with the HWE. Here, the HWE is expanded to account for polysomic inheritance, in which the alleles in a genotype are independent and randomly appear according to their frequencies. Thus, the genotypic frequency under the RCS can be expressed as

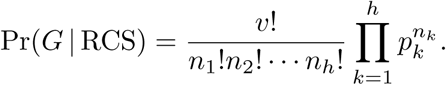

Where *p*_*k*_ denotes the frequency of *A*_*k*_. For example, the frequency of a tetraploid genotype *AABC* is 12*p*^2^_*A*_ *p*_*B*_*p*_*C*_.

When the conditions of the RCS are met, *α*_*i*_ = 0 whenever *i* > 0, then Equation (1) can be simplified to

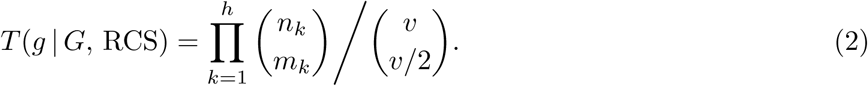

### Pure random chromatid segregation

Pure random chromatid segregation accounts for the crossing over and assumes the chromatids behave independently in meiotic anaphases, and are randomly segregated into gametes (Haldane, 1930, Figure 1B). Then the probability that a zygote *G* produces a gamete *g* at a multiallelic locus can be derived by sampling *v*/2 chromatids (or allele copies) in a total number of 2*v* chromatids (or allele copies). Because the number of chromatids is twice the number of chromosomes, the transitional probability *T* (*g* |*G,* PRCS) can be modified from Equation (2) by duplicating the terms *n*_*k*_ and *v* in those combinations as follows:

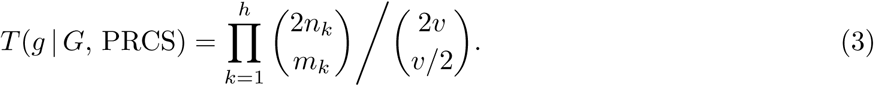

The value of alpha of PRCS can be derived by simulating the meiosis, first sampling *v*/2 − *i* IBDR allele pairs out of *v* IBDR allele pairs, second segregating *i* IBDR allele pairs and *v*/2 − 2*i* non-IBDR alleles into the gamete. Therefore, the value of *α*_*i*_ is given by

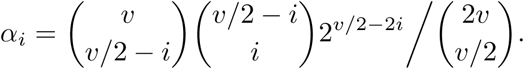

The values of alpha under PRCS, ranging from tetrasomic to dodecasomic inheritance are shown in Table 2. The expected number of IBDR allele pairs in a gamete *λ* = ∑ _*i*_ *α*_*i*_ = *v*(*v* − 2)/(16*v* − 8).

**Table 2:**
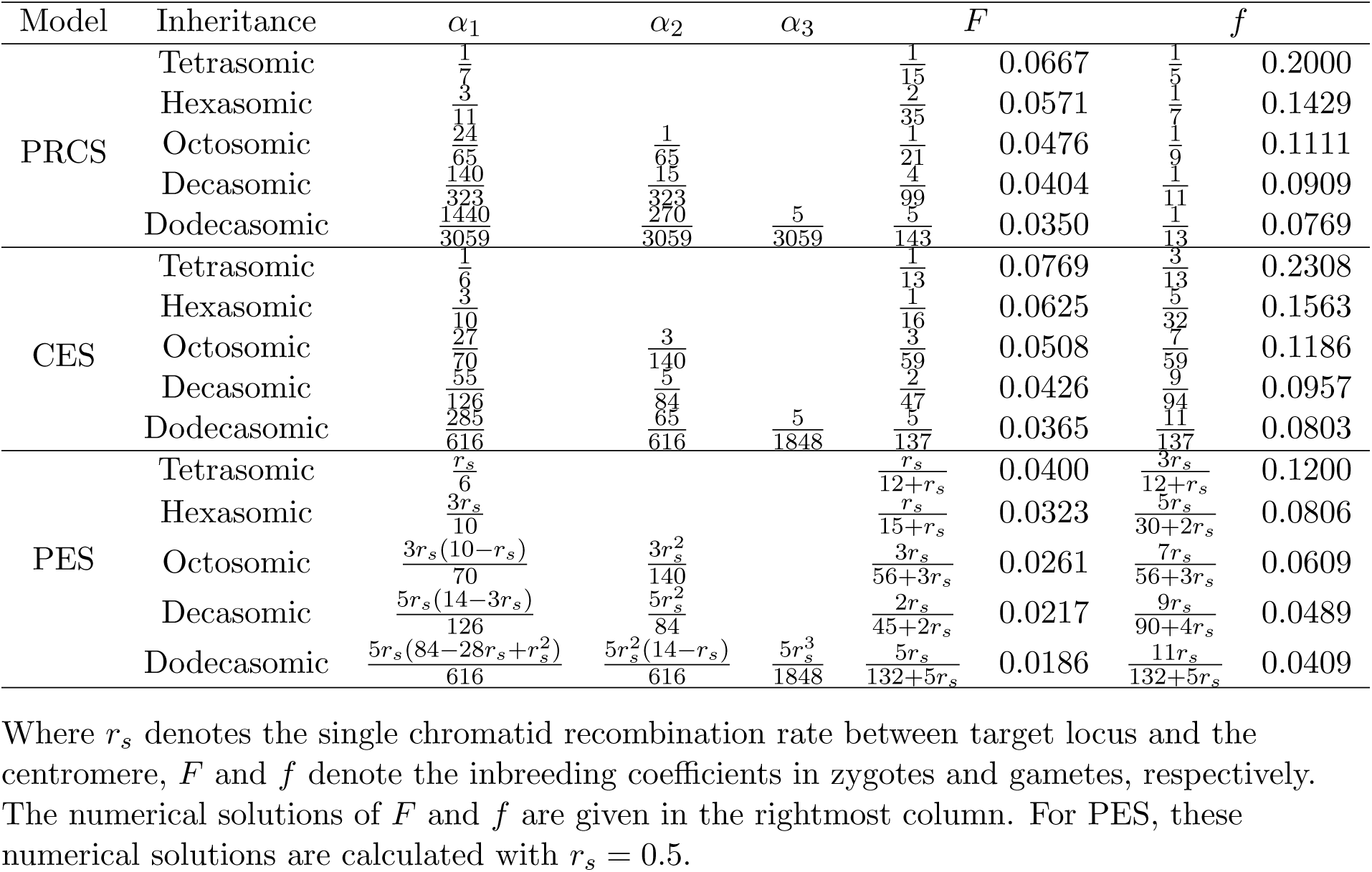
The probabilities of a gamete carrying *i* pairs of identical-by-double-reduction alleles (*α*_*i*_) and the inbreeding coefficient in an outcrossed population with pure random chromatid segregation (PRCS), complete equational segregation (CES) or partial equational segregation (PES) Where *r*_*s*_ denotes the single chromatid recombination rate between target locus and the centromere, *F* and *f* denote the inbreeding coefficients in zygotes and gametes, respectively. The numerical solutions of *F* and *f* are given in the rightmost column. For PES, these numerical solutions are calculated with *r*_*s*_= 0.5.

### Complete equational segregation

The *complete equational segregation* (CES) assumes the whole arms of the two pairing chromatids are exchanged between pairing chromosomes (Mather, 1935). In Figure 1C, the leftmost two chromosomes in the primary oocyte are paired in the long arms, as well as the rightmost two chromosomes. In Metaphase I, the chromosomes are randomly segregated into the secondary oocytes. If the pairing chromosomes are segregated in the same secondary oocyte, then the duplicated alleles may be further segregated into a single gamete.

The production of double-reduction gametes requires the fulfillment of two conditions: (i) the chromosomes paired at target locus are segregated into the same secondary oocyte; (ii) the duplicated alleles are segregated into the same gamete. In order to construct a mathematical model of CES, we simulate this meiosis, and the values of alpha will be derived by using a two-step process.

First, we model the probability, *P*_*j*_, that *j pairs of chromosomes paired at the target locus* (PCPs) are segregated into a secondary oocyte. There are *v*/2 PCPs in the primary oocyte, so we have 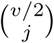 combinations to sample *j* pairs. Next, for the remaining *v*/2 − *j* PCPs, it still requires *v*/2 − 2*j* chromosomes that are not paired with each other at target locus during Prophase I to form the secondary oocyte. We therefore (i) sample *v*/2− 2*j* PCPs from the remaining *v*/2 − *j* PCPs (there are 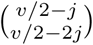 combinations); (ii) sample one chromosome from each PCP in the previously sampled *v*/2 − 2*j* PCPs (there are 2^*v*/2^−^2*j*^ chromosome combinations). The product of the above three numbers of combinations is divided by the total number of segregation modes in Metaphase I, i.e.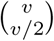, to derive *P*_*j*_. The process described above can be summarized by the following expression:

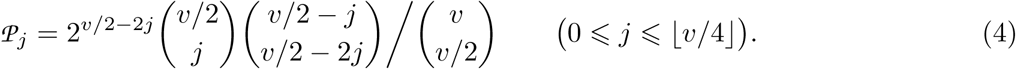

Second, we model the probability, *α*_*i*_, that *i* pairs of duplicated alleles are segregated into a single gamete. There are *j* PCPs in the second oocyte, and each has four alleles (e.g. the alleles in both chromosomes are *A* and *B* in the top second oocyte of Figure 1C). There are four segregation modes for each PCP in Anaphase II, where two of them (*AA* and *BB*) produce gametes with IBDR alleles and the other two (*AB* and *BA*) pass non-IBDR alleles into the gamete. In order to produce a gamete consisting of *i* pairs of IBDR alleles, *i* PCPs are sampled and segregate IBDR alleles. Following the description above, we have 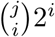 combinations of sampling *i* PCPs that pass IBDR alleles into a gamete from *j* PCPs. Next, the remaining *j* − *i* PCPs pass non-IBDR alleles, and so there are 2^*j*−*i*^ combinations. In each of the remaining *v*/2 − 2*j* chromosomes which are not paired with each other during Prophase I, one of the two alleles is segregated into the gamete, and hence there are 2^*v*/2^−^2*j*^ allele combinations. The product of these numbers of combinations is divided by the total number of segregation modes in Metaphase II, i.e. 2^*v*/2^, to obtain the expression 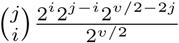, i.e.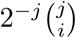, which is the value of *α*_i_ conditional on *j*, where *i*⩽ *j* ⩽ ⌊ *v*/4⌋. Because these events that the secondary oocyte contains different numbers of PCPs are mutually exclusive, their probabilities are weighted by *P*_*j*_ to obtain the weighted sum, i.e. *α*_*i*_, whose expression is as follows:

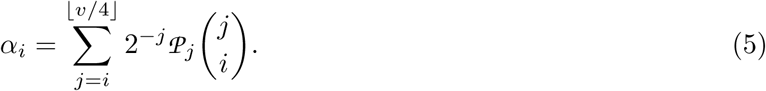

Using Equation (5), the values of alpha of CES can be derived for different inheritance modes, and these are presented in Table 2. The example deriving the values of alpha under octosomic inheritance is given in Appendix B. The expected number of IBDR allele pairs in a gamete *λ* = ∑_*i* _*α*_*i*_ = *v*(*v* − 2)/(16*v* − 16).

### Partial equational segregation

The *recombination fraction* (denoted by *r*) between two loci is defined as the ratio of the number of recombined gametes to the total number of gametes produced. The maximum recombination fraction for diploids is *r* = 0.5 (Xu, 2013). However, in autopolyploids, because there are *v*/2 PCPs, the maximum recombination fraction for an autopolyploid organism is *r* = 1 − 0.5^*v*/2^, e.g. *r* = 0.75 for autotetraploids.

We used the *single chromatid recombination rate* (denoted by *r*_*s*_) between the target locus and the centromere (*r*_*s*_) to evaluate the degree of recombination in polysomic inheritance, which is probability that an allele in a chromatid is exchanged with the pairing chromatid. Hence, *r* and *r*_*s*_ can be converted to each other by the formula *r* = 1 − (1 − *r*_*s*_)^*v*/2^. Moreover, *r*_*s*_ can be calculated by Haldane’s mapping function 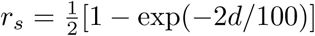 where d is the distance between the target locus and the centromere (the unit of length is the centimorgan).

In the CES model, the two alleles in a pair of pairing chromatids are assumed to be exchanged at a probability of 100%. This ‘ideal’ condition is unlikely to occur in nature. We thus usually consider that the maximum single chromatid recombination rate is *r*_*s*_ = 50%, and that the distance between the target locus and the centromere is extremely long. Although Mather (1935) incorporated the recombination rate into his model, assuming 100% recombination has been widely used in similar studies (e.g. Butruille and Boiteux, 2000; Wu, Rong Ling and Gallo-Meagher, Maria and Littell, Ramon C. and Zeng, Zhao Bang, 2001).

We incorporated the single chromatid recombination rate *r*_*s*_ into the CES to allow a greater generalization, which allows for the situation that only some fragments of the two pairing chromatids are exchanged between pairing chromosomes. Such a model is called the *partial equational segregation* (PES) model.

In the first step for CES, we model the probability *P*_*j*_ that *j* PCPs are segregated into a secondary oocyte, which remains unchanged in the PES.

Assuming that the alleles in each PCP are exchanged at a probability of *r*_*s*_ between pairing chromatids and those *j* PCPs are exchanged independently, then the number of *exchanged PCPs* (EPCPs) is drawn from the binomial distribution, and so the probability *𝒫*_*k*_ that the second oocyte contains *k* EPCPs is

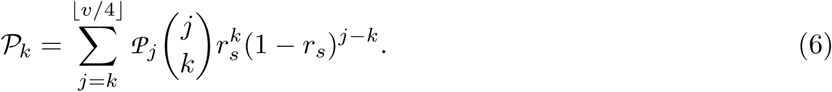

It is noteworthy that only the EPCPs are able to produce IBDR alleles under a PES model. Following the process of the second step for CES, we derive the alpha for PES by: (i) sampling *i* EPCPs from *k* EPCPs (there are 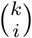ways to do this); (ii) sampling IBDR alleles from those *i* EPCPs (there are 2^*i*^ ways to do this); (iii) sampling non-IBDR alleles from the remaining *k* − *i* EPCPs (there are 2^*k*−*i*^ ways to do this); (iv) sampling the remaining non-IBDR alleles from the remaining *v*/2 − 2*k* chromosomes (there are 2^*v*/2^−^2*k*^ ways to do this). The product of the numbers of ways to do above four steps is now divided by the total number of segregation modes in Metaphase II, i.e. 2^*v*/2^, to obtain the expression 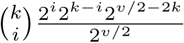, i.e 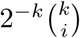, which is the value of *α*_*i*_ conditional on *k*, where *i*⩽ *k* ⩽ ⌊ *v*/4⌋. Moreover, the events that the secondary oocyte contains different numbers of EPCPs are mutually exclusive. Therefore the weighted sum of their probabilities with *𝒫*_*k*_ as their weights are calculated to obtain *α*_*i*_ under PES, whose expression is as follows:

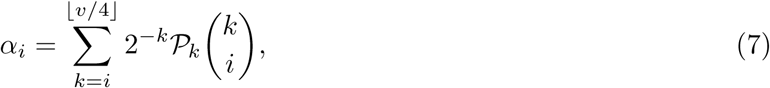

where *𝒫*_*k*_ is given in Equation (6).

Using Equation (7), the values of alpha in the PES model for tetrasomic to dodecasomic inheritance can be derived, and the results are presented in Table 2. The example deriving the values of alpha under octosomic inheritance is also given in Appendix B. The expected number of IBDR allele pairs in a gamete *λ* = ∑_*i*_ *α*_*i*_ = *r*_*s*_*v*(*v* − 2)/(16*v* − 16).

## DATA AVAILABILITY

We intend to archived our data in the Supporting Information and can be downloaded in the online page of this article.

**Material S1**: The appendices.

**Material S2**: The simulation computer program to obtain the numerical solutions of genotypic and phenotypic frequencies of zygotes and the genotypic frequencies of gametes. The program can be run on Windows platforms and requires the. Net Framework V2.0 runtime library.

**Material S3**: The C++ source code of the genotypic and phenotypic frequencies of polysomic inheritance under double-reduction. The prefix of functions is GFZ (genotypic frequencies of zygotes), GFG (genotypic frequencies of gametes), PFZ (phenotypic frequencies of zygotes), or PFG (phenotypic frequencies of gametes). An example that is the function in the second case (i.e. *G* = *AAAB*) in Equation (A4) is given by

~~~
double GFZ4_iiij(double a1, double pi, double pj)
{
 double pi2=pi*pi;
 return 8*(−1+a1)*pi2*(−3*a1+2*(−1+a1)*pi)*pj)/pow(2+a1,2);
}
~~~

## THE GENOTYPIC FREQUENCIES AT EQUILIBRIUM

Here we present two methods to derive the genotypic frequencies at a multiallelic locus for higher levels of ploidy. The first method, the non-linear method, is modified from Fisher’s (1943) method and uses the transitional probability from zygotes to gametes to account for the multiallelic loci. Because this method is computationally difficult for a ploidy level higher than six under current conditions, we develop an alternative method to derive the genotypic frequency up to and including decasomic inheritance. The second method, the linear method, uses the solution of the non-linear method at a biallelic locus as the initial solution, and at each step a novel allele is segregated from an existing allele and the genotypic frequencies are updated.

### Non-linear method

For this method, the total probability formula together with Equation (1) are used to derive the gamete frequencies and the multiplication theorem of probability is used to derive the zygote frequencies. If the genotypic frequencies reach equilibrium, they will not change subsequently, and the system of equations that we will construct can be solved by adding some constraints. In the following text, we denote GFG for *genotypic frequencies of gametes,* and GFZ for *genotypic frequencies of zygotes.* Our system of non-linear equations will be presented by the following two steps.

(i) Simulating meiosis. Given a gamete *g*, if the genotypic frequencies reach equilibrium, then the GFG is determined by the total probability formula, i.e. the sum of the transition probabilities *T* (*g* | *G* _*i*_) weighted by GFZ, symbolically

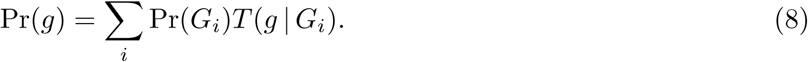

Here *G* _*i*_ is taken from all zygote genotypes and *T* (*g* | *G* _*i*_) can be obtained by Equation (1). Because there are 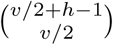 gamete genotypes, Equation (8) represents 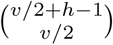 equations.

(ii) Simulating fertilization. The GFZ in the next generation is determined by the multiplication theorem of probability with GFG as the weight. Under equilibrium, the GFG and GFZ are constant across generations, then for a zygote *G,* we can establish an equation as follows:

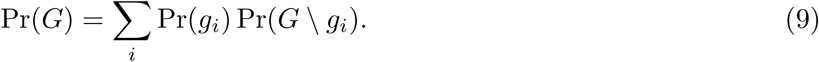

Where *G* is regarded as a multiset; *g*_*i*_ is taken from all possible different subsets of *G* with *v*/2 elements; and the symbol \ denotes the operation of set difference. Here, a *multiset* is a generalized set, whose elements can be repeated (e.g. if *G* = *AAAB*), it can be regarded as a multiset consisting of four elements, i.e. *G* = {*A, A, A, B*}, and its different subsets containing two elements are {*A, A*} and {*A, B*}; moreover, if *g* = *AA* is regarded as *g* = {*A, A*}, then *G* \ *g* = {*A, B*}. Because there are 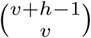 zygote genotypes, Equation (9) represents 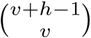 equations. If the double-reduction ratios in two parents are different, then the genotypic frequencies of egg and sperm should be treated separately in Equation (9), e.g. Pr(*g*_*i*_) and Pr(*G* \ *g*_*i*_) are replaced by the frequencies of an egg *g*_*i*_ and a sperm *G* \ *g*_*i*_, respectively, and their frequencies are calculated by substituting different double-reduction parameters into Equation (8).

The process that transforms the allele frequencies into the GFG can be described by a linear substitution in the sense that every allele frequency can be expressed as a linear combination of the GFG. Substituting the equations determined by Equation (9) into the equations determined by Equation (8), we obtain a system of non-linear equations with the GFG as unknowns. This system of equations together with those expressions of the linear combinations mentioned above still forms a system of non-linear equations with the GFG as unknowns and the allele frequencies as parameters. For a lower level of ploidy, the latter system of non-linear equations can be solved, whose solution is with parameters. An example is given in Appendix C, which shows how to find the solution of the latter system of non-linear equations (i.e. how to derive the expressions of GFG with parameters) for tetrasomic inheritance at a triallelic locus.

In Appendix C, for the case of GFG, the whole gamete genotypes are *AA, BB, CC* and *AB, AC, BC*. We choose *AA* and *AB* as representatives of genotype pattern, then each genotype can be classified into one of these patterns. The partial solution determined by the representatives is called the *generalized form* of the solution mentioned above, whose expression is given in Equation (A3). For the case of GFZ, the notion of generalized form can be similarly defined, whose expression is given in Equation (A4).

For each gamete with *v*/2 allele copies, the number of alleles *h* at the target locus should at least be *v*/2 to allow for all genotypic patterns to be displayed. For example, the full gamete heterozygote *ABCD* under octosomic inheritance can be observed at a tetra-allelic locus. Further, if *h* ⩾ *v*/2 + 1, the generalized form for all genotypic patterns of GFG can be obtained. Because the allele frequencies at each locus sum to unity, the number of degrees of freedom is equal to *h* − 1. The full gamete heterozygotes can therefore be expressed by *h* − 1 allele frequencies.

Using the tetrasomic inheritance under RCS as an example, the frequency of a gamete *g* = *AB* can be solved at a biallelic locus, and the solution is 2*p*_*A*_(1 - *p*_*A*_). Moreover, its solution with the generalized form at a triallelic locus can also be obtained, with the answer being 2*p*_*A*_*p*_*B*_.

The number of equations determined by Equation (8) or Equation (9) increases exponentially with increasing levels of ploidy. For example, for the case of Equation (8), the number equals 6, 20, 70, 252 and 924 for tetrasomic inheritance to dodecasomic inheritance, while the corresponding sequence is 15, 84, 495, 3003 and 18564 for the case of Equation (9). The two numerical sequences increase by about and 6.0 times for each level, respectively. Moreover, each expression also becomes more complex as the ploidy level increases. For the case of dodecasomic inheritance, it takes multiple gigabytes of space to store these expressions.

Therefore the system of non-linear equations becomes computationally difficult for a higher level of ploidy. Using this method, we solved the GFG and GFZ for a maximum ploidy level of six on a workstation with two Intel Xeon E5 2696 V3 CPUs, which have 36 cores in total.

### Linear method

An alternative method to solve the genotypic frequencies at equilibrium is via the linear method. For a working example we focus on a protein-coding gene in a panmictic population at equilibrium with an infinite number of individuals. Suppose that there are three alleles with a unique DNA sequence, denoted by *A, B* and *C*, and let *p*_*A*_, *p*_*B*_ and *p*_*C*_ be their frequencies. Due to the degeneration of codons, two alleles (e.g. *B* and *C*) may code for the same protein. If isozyme electrophoresis is used for genotyping, then these two alleles cannot be distinguished, so both of them will, for example, be typed as *B*. However, if DNA sequencing is used, both alleles can be distinguished.

Therefore, it can be inferred that the number of alleles of this gene depends on how it is typed, with both biallelic and triallelic states conforming to equilibrium assumptions (i.e. large population size, random mating, no mutation, no selection and no migration) and genotypic frequencies should concur with the equilibrium.

This scenario can also be expressed as follows: (i) originally, the locus is biallelic and the distribution of genotypes is at equilibrium; (ii) at each step, one novel allele mutates from an existing allele, the mutation rate is constant among different copies of this allele in all genotypes, and the IBD alleles in a genotype mutate simultaneously; (iii) therefore, after the mutation at each step, the genotypic frequency will still concur with the equilibrium; (iv) the mutations are repeated until there are *v*/2 + 1 alleles to obtain the generalized form of GFG. The following deductions are therefore derived.

**Deduction (i).** For both of GFZ and GFG, the frequency of a genotype that contains only the unchanged alleles remains unchanged. In fact, the analytical expression of genotypic frequencies can be regarded as a function of the allele frequencies (refer to Equations (A3) and (A4) in Appendix C). Therefore, when a genotype contains only the unchanged alleles, the corresponding expression of the function is invariable in the process of mutation.

**Deduction (ii).** For both of GFZ and GFG, the summations of the frequencies before and after the mutation for each genotype that has the same composition of unchanged alleles are equal (e.g. *P*_*AABB,2*_ = *P*_*AABB,3*_ + *P*_*AABC,3*_ + *P*_*AACC,3*_ where the last digit in every subscript denotes the number of alleles observed). This fact can be interpreted as several genotypes being merged into one when the locus switches from a triallelic state to a biallelic state.

**Deduction (iii).** For both of GFZ and GFG, the ratio of frequencies of changed alleles in the genotypes that have the same composition of unchanged alleles is equal to the ratio of frequencies of changed alleles in the population. For example, if the changed alleles are *B* and *C*, we have *P*_*AAAB,3*_ : *P*_*AAAC,3*_= *p*_*B,3*_ : *p*_*C,3*_ and 2*P*_*AABB,3*_ + *P*_*AABC,3*_ : 2*P*_*AACC,3*_ + *P*_*AABC,3*_ = *p*_*B,3*_ : *p*_*C,3*_. Because the mutation rate is constant in different copies of *B* in all genotypes, a fixed proportion 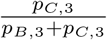 of *B* are mutated to *C*.

**Deduction (iv).** For two genotypes with the same pattern, we can use the expression of one genotype frequency to derive that of the other. For example, if the analytical expression *P*_*AAAB,3*_ = *f* (*p*_*A,3*_, *p*_*B,3*_) of the function is given, by replacing *p*_*A,3*_ with *p*_*B,3*_, and *p*_*B,3*_ with *p*_*C,3*_, it follows the expression of *P*_*BBBC,3*_, that is *P*_*BBBC,3*_ = *f* (*p*_*B,3*_, *p*_*C,3*_).

According Deduction (iv), if a generalized form for the GFG or GFZ is given, we can write down the whole expressions of the GFG or GFZ.

Using these deductions as stated above, the GFG can be solved step-by-step. At each step, one novel allele is separated from an existing allele. The procedure begins with a biallelic locus, which can be solved by the non-linear method. After the number of alleles reaches *v*/2 + 1, the generalized form of GFG is obtained, and then the GFZ can be derived by using Equation (9).

Using the hexasomic inheritance as an example, we assume that the GFG at a biallelic locus is solved by the non-linear method, i.e. the expressions of *P*_*AAA,2*_, *P*_*AAB,2*_ and *P*_*ABB,2*_ are derived. If a new allele (e.g. *C*) is segregated from *B*, we obtain *P*_*AAA,2*_ = *P*_*AAA,2*_ via Deduction (i), and 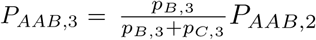 via Deduction (iii). Additionally, *P*_*BBB,3*_ and *P*_*CCC,3*_ can be derived from *P*_*AAA,3*_ via Deduction (iv). Similarly, *P*_*AAC*,3_, *P*_*ABB,3*_, *P*_*BBC,3*_, *P*_*ACC,3*_ and *P*_*BCC,3*_ can be derived from *P*_*AAB,3*_. Furthermore, *P*_*ABC,3*_ will be obtained from *P*_*ABB,2*_ - *P*_*ACC,3*_ - *P*_*ABB,3*_ via Deduction (ii). Because the degrees of freedom are currently two, one more step is required. In this step, *P*_*ABC*_ can be updated by Deduction (iii), and expressed by 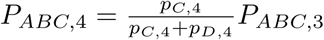. Finally, *P*_*AAA*_ and *P*_*AAB*_ remain unchanged because of Deduction (i), and the remaining GFG will be obtained via Deduction (iv). The results of GFG and GFZ for hexasomic inheritance are presented in Appendix D.

The linear method can also be characterized by a system **Ax** = **b** of linear equations. Using the first mutation in hexasomic inheritance as an example, the coefficient matrix **A** is established (see Table 3 for details, in which *p, q* and *r* denote *p*_*A,3*_, *p*_*B,3*_ and *p*_*C,3*_, respectively). The elements in **b** can be derived from the above four deductions. For example, the first seven equations of **Ax** = **b** are

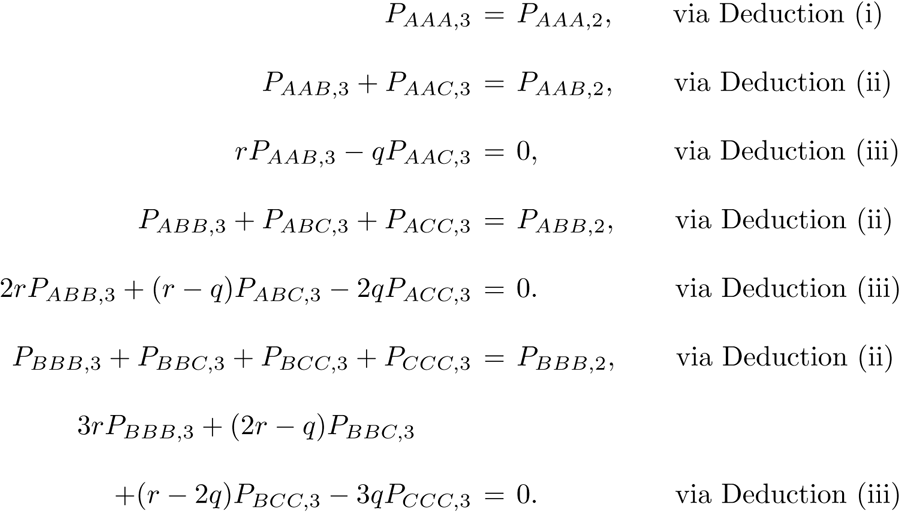

Then the first seven elements in **b** are then *P*_*AAA,2*_, *P*_*AAB,2*_, 0, *P*_*ABB,2*_, 0, *P*_*BBB,2*_, 0. For these seven equations, one is derived by Deduction (i), where one is number of unchanged genotypes; three are derived by Deductions (ii) and (iii), respectively, where 3 is the number of changed genotypes. Therefore, with the first three deductions, we are able to establish the core equations. Further, by Deduction (iv) we can write down the remaining equations and the total number of equations is *h* times of the number of core equations.

**Table 3:** Matrix **A** of the linear method of hexasomic inheritance at triallelic locus Where *p, q* and *r* denote the frequencies of alleles *A, B* and *C* at a triallelic locus, respectively. The blank elements are zero.

| Table 3: Matrix **A** of the linear method of hexasomic inheritance at triallelic locus |  |  |  |  |  |  |  |  |  |
| --- | --- | --- | --- | --- | --- | --- | --- | --- | --- |
| Genotypic frequencies of gamete |  |  |  |  |  |  |  |  |  |
| $P_{AAA,3}$ | $P_{AAB,3}$ | $P_{AAC,3}$ | $P_{ABB,3}$ | $P_{ABC,3}$ | $P_{ACC,3}$ | $P_{BBB,3}$ | $P_{BBC,3}$ | $P_{BCC,3}$ | $P_{CCC,3}$ |
| 1 |  |  |  |  |  |  |  |  |  |
|  | 1 | 1 |  |  |  |  |  |  |  |
| | $r$ | $-q$ | | | | | | | |
|  |  |  | 1 | 1 | 1 |  |  |  |  |
| | | | $2r$ | $r - q$ | $-2q$ | | | | |
|  |  |  |  |  |  | 1 | 1 | 1 | 1 |
| | | | | | | $3r$ | $2r - q$ | $r - 2q$ | $-3q$ |
|  |  |  |  |  |  | 1 |  |  |  |
|  |  |  | 1 |  |  |  | 1 |  |  |
| | | | $r$ | | | | $-p$ | | |
|  | 1 |  |  | 1 |  |  |  | 1 |  |
| | $2r$ | | | $r - p$ | | | | $-2p$ | |
| 1 |  | 1 |  |  | 1 |  |  |  | 1 |
| $3r$ | | $2r - p$ | | | $r - 2p$ | | | | $-3p$ |
|  |  |  |  |  |  |  |  |  | 1 |
|  |  |  |  |  | 1 |  |  | 1 |  |
| | | | | | $q$ | | | $-p$ | |
|  |  | 1 |  | 1 |  |  | 1 |  |  |
| | | $2q$ | | $q - p$ | | | $-2p$ | | |
| 1 | 1 |  | 1 |  |  | 1 |  |  |  |
| $3q$ | $2q - p$ | | $q - 2p$ | | | $-3p$ | | | |

The numbers of rows and columns of **A** are 21 and 10, respectively, which means that there are 21 equations and 10 unknowns (i.e. the whole GFG after mutation, e.g. *P*_*AAA,3*_ and *P*_*AAB,3*_ and so on.) in this system **Ax** = **b**. There are thus many non-independent equations. The rank of **A** can be derived by symbolic calculation, whose value is 10, equal to the number of unknowns. Therefore our system of linear equations has a unique solution: **x** = (**A**^*T*^ **A**)^-1^**A**^*T*^ **b**.

The numbers of rows and columns of **A** are respectively 39 and 28 for dodecasomic inheritance at a triallelic locus, and the rank of **A** is 27 by symbolic calculation. Unfortunately, because there are 28 unknowns, our system **Ax** = **b** is underdetermined and has an infinite number of solutions.

Using this method, we are able to derive the GFZ and GFG from tetrasomic to decasomic inheritance. For the cases of tetrasomic and hexasomic inheritance, these results are identical to the solutions obtained from the non-linear method. Subsequent expressions become more complex with increasing levels of ploidy, so we only present the results for hexasomic inheritance in Appendix D. As an alternative, these expressions are placed in a 660 KB C++ file, and the source code is available in our supplementary material.

Due to the genotypic ambiguity of polyploid species, the correct genotype cannot be obtained because some genotypes share a common electrophoretic band type in PCR-based co-dominant markers (e.g. microsatellites) and the allelic dosage is unknown (Hardy, 2016). For example, in autotetraploids, the three genotypes *AAAB, AABB* and *ABBB* have a same electrophoretic band type of *AB*. We define the set of alleles within an individual as its phenotype, denoted by *G′*. The phenotypic frequencies can then be calculated by taking the summation of the frequencies of all possible genotypes that yield this phenotype, and they are also given in the source code in our supplementary material.

### Inbreeding coefficient and heterozygosity

Double-reduction has a similar effect as inbreeding, both not only resulting in IBD alleles in the zygote but also slightly increasing homozygosity (Hardy, 2016). Here, we assess the impact of inbreeding and double-reduction on the inbreeding coefficient. The concept of inbreeding is relative to a reference population, which by definition is without inbreeding or relatedness, with all alleles in the reference population defined as not being IBD (Lynch and Walsh, 1998).

The *inbreeding coefficient* in polysomic inheritance is defined as the probability of sampling two IBD alleles from a genotype without replacement (Huang *et al*., 2015). In polyploids, gametes also have multiple alleles at a target locus, and the inbreeding coefficient and heterozygosity can also be applied to gametes to measure the degree of IBD or *identical-by-state* (IBS) allele pairs (Barone *et al*., 1995).

After fertilization, there are 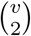 pairs of alleles in a zygote, where 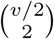 pairs of alleles are from one gamete, and *v* ^2^/4 pairs of alleles are from different gametes. The numbers of IBD allele pairs in both types of allele pairs are derived as follows.

The number of IBDR allele pairs in the 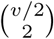 allele pairs from the same gamete is the expected value *λ* of allele pairs, where 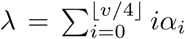. In the remaining allele pairs, the number of IBD allele pairs inherited from the parent is a proportion *F* of those remaining, where *F* is the inbreeding coefficient in zygotes of the previous generation. Therefore, there are 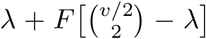 pairs of IBD alleles in this gamete. Hence the inbreeding coefficient for this gamete is as follows:

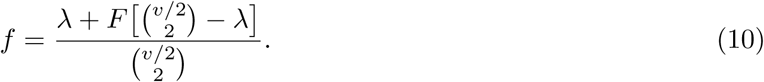

For the *v*^2^/4 allele pairs from two different fertilizing gametes, the mating individuals in the presence of inbreeding may also share IBD alleles. The co-ancestry coefficient is used to measure the degree of kinship in mating individuals, where the *co-ancestry coefficient*, denoted by *θ*, is defined as the probability that two alleles, one randomly sampled from each individual, are IBD (Jacquard, 1972). Then there is a value *θv*^2^/4 expected IBD allele pairs between fertilizing gametes.

From the above derivation, the number of pairs of IBD alleles in the zygote is in total 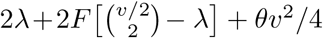. Therefore, the inbreeding coefficient for zygotes of the current generation *F*′ is as follows:

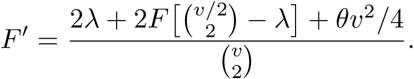

The formula is the transition function of inbreeding coefficients. If the genotypic frequencies reach equilibrium, these inbreeding coefficients will be constant across generations. By replacing *F*′ with *F* in the transition function, the inbreeding coefficient *F* at equilibrium can be solved, whose expression is as follows:

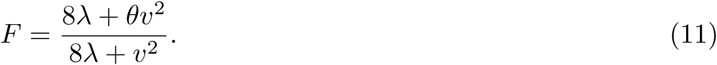

For an outcrossed population, *θ* = 0. Then, by substituting Equation (11) for Equation (10), the inbreeding coefficients *F* and *f* can be obtained, whose values at equilibrium for different double-reduction models are shown in the rightmost two columns of Table 2.

If the relationships between mating individuals can be classified into several types (e.g. self: selfing, parent-offspring: backcross), then the co-ancestry coefficient *θ* can be derived by a weighted average of the co-ancestry coefficients of those types of relationships. An example to derive the expressions of these co-ancestry coefficients as well as the inbreeding coefficient *F* is given in Appendix E.

Following the modification of the definition of inbreeding coefficients, the *heterozygosity* in polysomic inheritance is defined as the probability of sampling two non-IBS alleles from a genotype without replacement (Hardy, 2016). For the two alleles sampled, if they are IBD, then these alleles cannot be IBS; otherwise they are independent and are non-IBS alleles at a probability of 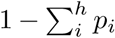. Let *H* be the heterozygosity of zygotes and *h* be that of gametes, then:

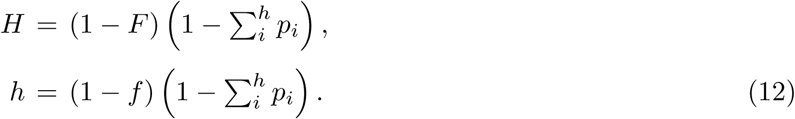

Where *F* is the inbreeding coefficient of zygotes and *f* is that of gametes. These expressions are identical to Nei’s (1977) estimator of inbreeding coefficients, i.e.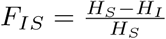, where *H*_*I*_ and *H*_*s*_ are the observed and expected heterozygosities, respectively.

### Simulation

The numerical solutions of genotypic and phenotypic frequencies of gametes and zygotes calculated by computer simulations will be used to compare with the analytical solutions so as to validate our models.

A population is generated with its initial genotypic frequency drawn from HWE. Because we do not store the specific genotype for each individual, but only the frequency of each genotype, the population size can be considered as infinite. To display all kinds of genotype patterns, the number of alleles *h* should be greater than or equal to the ploidy level. For example, the complete heterozygote in tetrasomic inheritance *ABCD* cannot be observed at a triallelic locus. For simplicity, suppose that the allele frequencies are uniformly distributed, i.e. the frequency of any allele is equal to 1/*h*, so that the frequencies of genotypes or phenotypes with a same pattern are equal (e.g. *P*_*AAAB*_ = *P*_*BBBC*_).

The double-reduction models are simulated to calculate the gamete frequencies. First, for PRCS, the alleles in the zygotes are duplicated and then are randomly segregated into four gametes. Second, for CES, the chromosomes are randomly paired and the alleles are exchanged between pairing chromosomes, and then the chromosomes are randomly segregated into two secondary oocytes, finally the alleles within the same chromosomes are randomly segregated into two gametes. Third, for PES, the chromosomes are randomly paired and the alleles are exchanged between paired chromosomes at a probability of *r*_*s*_, and the remaining steps are the same as CES.

For a zygote, there are many possible ways to generate gametes. We enumerate each possibility and calculate the weighted average of gamete frequencies according to the probability of each possibility. The frequencies of identical gametes produced from different zygotes are weighed again by the zygote frequencies so as to obtain the gamete frequencies in the population.

The gametes are randomly merged to simulate fertilization, then the zygote frequency in the next generation can be calculated. The population is reproduced for 100 generations to converge the genotypic frequencies. The simulation program can be found in the electronic supplementary materials.

There are many potential genotypes and phenotypes in a population, and it is thus impractical to show all of these frequencies in the manuscript. As an alternative, the phenotypic frequencies of zygote patterns are shown in Table 4, and the genotypic frequencies of gamete patterns are shown in Table 5. As a comparison, the corresponding analytical solutions of model predictions are also shown. It is clear from the data shown in Tables 4 and 5 that both of numerical and analytical solutions of phenotypic frequencies of zygotes are approximately equal, so are the genotypic frequencies of gametes obtained via different approaches.

**Table 4:**
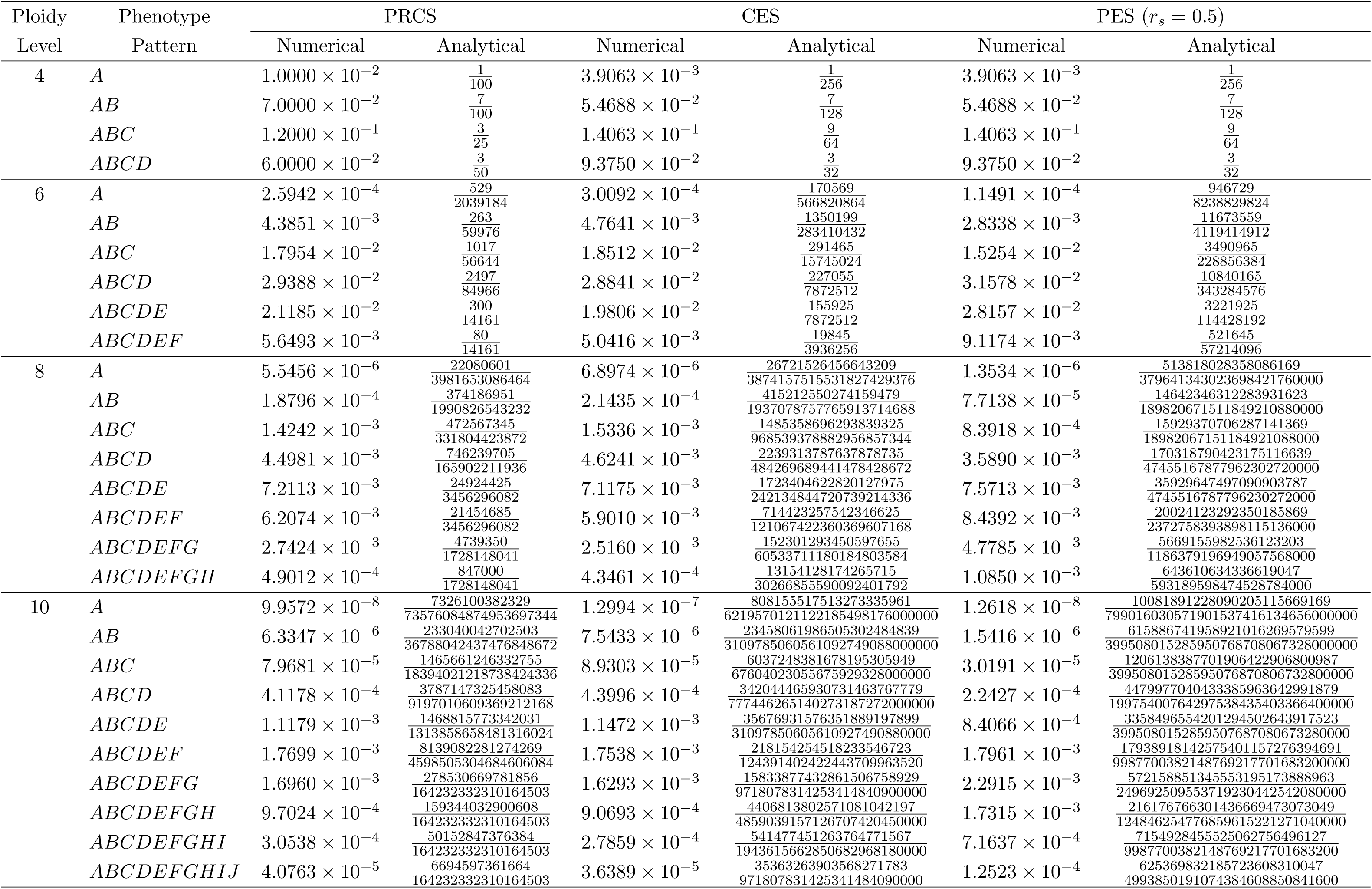
The numerical and analytical solutions of phenotypic frequencies obtained by numerical simulation and model predication. The number of alleles at this locus is equal to the ploidy level and their frequencies are uniformly distributed.

**Table 5:**
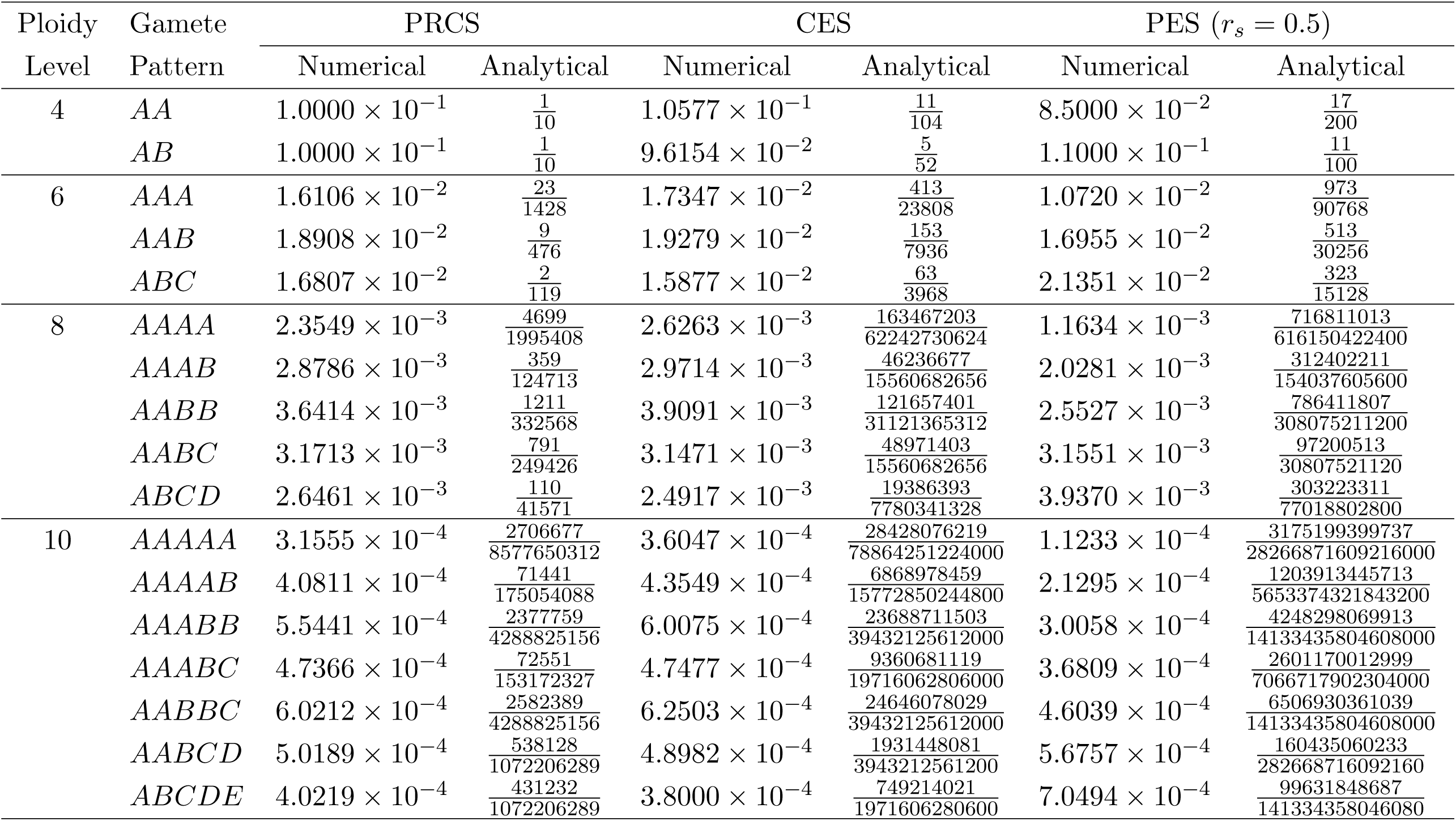
The numerical and analytical solutions of gamete frequencies obtained by numerical simulation and model predication. The number of alleles at this locus is equal to the ploidy level and their frequencies are uniformly distributed.

## DISCUSSION

### Double-reduction models

Here, we describe existing double-reductions models and extend them to the systems that account for any even level of ploidy. However, some of these models assume ideal situations that are unrealistic in nature. The CES model assumes that the single chromatid recombination rate *r*_*s*_ is one at any locus, but the maximum value of *r*_*s*_ is 50% if the locus is located extremely far from the centromere. We incorporate the single chromatid recombination rate into CES, and define this as the partial equational segregation (PES).

Although PES probably reflects more closely to natural patterns of occurrence and may thus more accurately explain real genotypic frequencies, other models (RCS, PRCS and CES) are still useful. Researchers are able to apply them to calculate the genotypic frequencies in their applications without estimating alpha. In contrast, the PES needs an additional parameter *r*_*s*_ to calculate the genotypic frequencies, and thus it has one more degree of freedom if *r*_*s*_ is estimated from the same data. Although PES may perform better in calculating the likelihood from genotypic/phenotypic data, it is also penalized more via the AIC (Akaike information criterion, Akaike, 1974) or the BIC (Bayesian information criterion, Schwarz *et al.*, 1978).

In addition, all of these models (RCS, PRCS, CES and PES) assume that the chromosomes must form multivalents during Prophase I. However, meiosis of polyploids can be much more complex. Some allopolyploid species display a mode of inheritance intermediate between disomic and polysomic at some loci (segmental allopolyploids, Stift *et al.*, 2008), and some autopolyploid species also form bivalent, univalent and other types of valent during meiosis (Qu *et al.*, 1998; Lloyd and Bomblies, 2016). The formation of different types of valent may influence the sterility of the gametes or seeds (Swaminathan and Sulbha, 1959; Crowley and Rees, 1968; Solís Neffa and Fernández, 2000). Such a model can hardly be established if we consider so many factors, e.g. fertility, proportion and double-reduction ratio for each kind of valent.

Regardless of how complex the nature of meiosis, different models will yield the same result: the IBDR alleles are in the fertile gamete. Therefore, all of these models can be incorporated into a generalized framework, i.e. using the values of alpha to express a model. Meanwhile, comparing with RCS, PRCS and CES models, the number of degrees of freedom of such a model increases only by ⌊*v*/4⌋.

### Genotypic frequencies, inbreeding coefficient and heterozygosity

Two methods are proposed to solve the genotypic equilibrium under double-reduction at a locus with an arbitrary number of alleles. The first method is the non-linear method, which can theoretically be applied to any even level of ploidy. However, it becomes computationally difficult for organisms with high levels of ploidy because these organisms have too many potential genotypes (e.g. there are 19494 nonlinear equations in dodecasomic inheritance). Although the system of non-linear equations for a higher level of ploidy is computationally difficult and we can currently only solve the genotypic frequencies for hexasomic inheritance, it will be possible with more powerful computers and more advanced algorithms.

The second method is the linear method, which requires more steps but less computational power. This uses the solution obtained by the non-linear method at a biallelic locus as the initial solution, and at each step it is assumed that one novel allele mutates from an existing allele. Some novel genotypes will be separated from existing genotypes, whilst the genotypic frequencies will still concur with the equilibrium. Using the four deductions, the equations of the genotypic frequencies of gametes are initially established by the total probability formula, and then the equations of the genotypic frequencies of zygotes are established by the multiplication theorem of probability. Unfortunately, the constraints are insufficient to obtain a unique solution for ploidy levels greater than or equal to 12. Although this is not perfect, our method is more than adequate to answer most current research questions on this subject.

Double-reduction results in not only in gametes carrying IBD alleles but also slightly increases homozygosity (Hardy, 2016). It has similar effects as inbreeding, and can cause a deviation in genotypic frequencies from the Hardy-Weinberg equilibrium (Luo *et al.*, 2006). By modifying the definition of the inbreeding coefficient and heterozygosity in polysomic inheritance, the symbolic expressions measuring how double-reduction and inbreeding influence the inbreeding coefficient and heterozygosity are derived. We found that double-reduction can cause a maximum inbreeding coefficient being 0.0769 (in CES) under tetrasomic inheritance in an outcrossed population. For the same double-reduction model, the double-reduction ratio is increased whilst the inbreeding coefficient is reduced with increasing levels of ploidy (Table 2).

In the presence of double-reduction and/or inbreeding, the theoretical observed heterozygosity can be calculated from the inbreeding coefficient, and is identical to Nei’s (1977) estimator of the inbreeding coefficient. This means that the inbreeding coefficient in polysomic inheritance can still be estimated by previous estimators, the same as for other *F* - statistics, although some modifications should be made to account for the finite sample size (e.g. Robertson and Hill, 1984; Weir and Cockerham, 1984).

### Applications of zygote frequencies

Genotypic frequencies can be used for many applications, such as for population genetics, molecular ecology, molecular breeding, and so on. Some of the applications of zygote frequencies are listed as follows.

(i) Estimating the allele frequency from phenotypes: the probability that observing a genotype conditional on a phenotype can be derived from the GFZ and Bayes formula. By counting the number of copies of each allele in each possible genotype and using the conditional probability as a weight, the allele frequencies can be updated from an initial solution with an Expectation Maximization algorithm (e.g. de Silva *et al.*, 2005; Kalinowski and Taper, 2006).
(ii) Based on the allele frequency, many subsequent analyses can be performed. For example, genetic diversity analysis (e.g. Rousset, 2008), genetic distance analysis (e.g. Nei, 1972), population differentiation analysis (e.g. Cockerham, 1973), analysis of molecular variances (Excoffier and Lischer, 2010), principal coordination analysis (e.g. Peakall and Smouse, 2012), and hierarchy clustering (e.g. Odong *et al.*, 2011).
(iii) Population assignment and Bayesian clustering: by calculating the product of multilocus genotypic frequencies, the probability of randomly sampling an individual with certain multilocus genotypes from a population can be obtained (i.e. likelihood). The individual is assigned to the population with maximum likelihood (e.g. Peakall and Smouse, 2012). In the non-admixture model of Bayesian clustering, the population origin of each individual is randomly drawn from the populations according to the posterior probability that the individual originated from each population, and is updated iteratively in a Markov Chain Monte Carlo (MCMC) algorithm (Pritchard *et al.*, 2000). The posterior probability is calculated from the multilocus genotypic frequencies of the individual from each population by the Bayes formula (Pritchard *et al.*, 2000).
(iv) Equilibrium test: as a Hardy-Weinberg equilibrium test, this test is able to evaluate whether the distribution of genotypes/phenotypes concurs with the equilibrium state for a particular double-reduction model. The observed number of each genotype/phenotype can be obtained by counts from the data, and the expected number can be obtained by the GFZ or PFZ. A Chi-square goodness-of-fit test or a Fisher’s *G* test can be used for this application (e.g. Rousset, 2008).
(v) Linkage disequilibrium test: the contingency table of a pair of loci is established, with each row denoting a genotype of the first locus with each column denoting that of the second locus. The expected number in each cell under the hypothesis that the two loci are under linkage equilibrium is the product of the GFZ at these two loci (e.g. Rousset, 2008).
(vi) Estimating individual inbreeding coefficient, relatedness coefficient or kinship coefficient from phenotypes: method-of-moment estimators (e.g. Hardy and Vekemans, 2002; Huang *et al.*, 2014) or the maximum-likelihood estimator (e.g. Huang *et al.*, 2015) can be used to estimate each of these coefficients between possible genotypes, with the estimates being weighted according to the conditional probability in application (i) above.

### Applications of gamete frequencies

There are also some potential applications of the transitional probability from zygotes to gametes and the gamete frequencies, in additional to solve the values of alpha for PRCS (Equation (3)) and to derive the genotypic frequencies at equilibrium (Equation (8)).

(i) Parentage analysis: the transitional probability that a parent or a pair of parents produce an offspring is used in the calculation of likelihoods of the following two hypotheses: (i) is that an alleged father is the true father of the offspring; (ii) is that the alleged father is unrelated to the offspring and is randomly sampled from the population (Marshall *et al.*, 1998; Kalinowski *et al.*, 2007). This transitional probability is a summation of the products of two gamete frequencies that form the offspring’s genotype, where the frequency is equal to Pr(*g* | *G*_*a*_) Pr(*G*_*o*_ \ *g* | *G*_*m*_) if the mother’s genotype is known, or equal to Pr(*g* | *G*_*a*_) Pr(*G*_*o*_ \ *g*) if the mother’s genotype is unknown, in which *G*_*o*_, *G*_*a*_ and *G*_*m*_ denote the genotypes of the offspring, alleged father and true mother, respectively.
(ii) Deriving the segregation ratio in mapping populations, or deriving the genotypic frequencies for populations which are not at equilibrium: we use an F_2_ population in tetrasomic inheritance as an example. Assuming the genotypes of F_1_ individuals are *AABB*, using Equation (1), the gamete ratio is *AA* : *AB* : *BB* = 1 + 2*α*_1_ : 4 − 4*α*_1_ : 1 + 2*α* _1_. From the multiplication theorem, the segregation ratio of the F_2_ population is 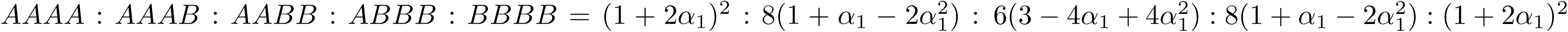.
(iii) Estimating the values of alpha from mapping populations: by equating the observed segregation ratio with the expected value, a system of non-linear equations can be established. The values of alpha can be solved by the least-squares method.
(iv) Perform an equilibrium test or a linkage disequilibrium test: the deviation of the observed gamete frequencies from expected can be measured by either a *χ*^2^ or a *G* statistic by using either a Chisquare test or a Fisher’s *G*-test, respectively. With the cumulative distribution function of a Chi-squared distribution, the significance can be calculated.

## ACKNOWLEDGEMENTS

This study was funded by the Strategic Priority Research Program of the Chinese Academy of Sciences (XDB31020302), the National Natural Science Foundation of China (31770411, 31730104 and 31770425), and the Young Elite Scientists Sponsorship Program by CAST (2017QNRC001), the National Key Programme of Research and Development, the Ministry of Science and Technology of China (2016YFC0503202), and the Natural Science Basic Research Plan in Shaanxi Province of China (2018JM3024). DWD is supported by a Shaanxi Province Talents 100 Fellowship.

## AUTHOR CONTRIBUTIONS

BGL & KH designed the research, KH and TCW wrote the program and draft, XXC and RCL performed simulations, PZ checked the data, and DD modified the manuscript.

## MATERIAL S1: THE APPENDICES

### A Derivation of Pr(*BBBB* | *ABBBBBBB*)

In this example, we show how to derive the transitional probability from a zygote genotype *G* = *ABBBBBBB* to a gamete genotype *g* = *BBBB*. According to Equation (1), *m*_*k*_ and *n*_*k*_ are respectively the numbers of copies of *A*_*k*_ in *g* and *G, h* is the number of alleles at this locus, and *v* is the ploidy level. Letting *v* = 8, *h* = 2, *m*_1_ = 0, *m*_2_ = 4, *n*_1_ = 1 and *n*_2_ = 7, and expanding the sum formula in Equation (1), it follows the following expression:

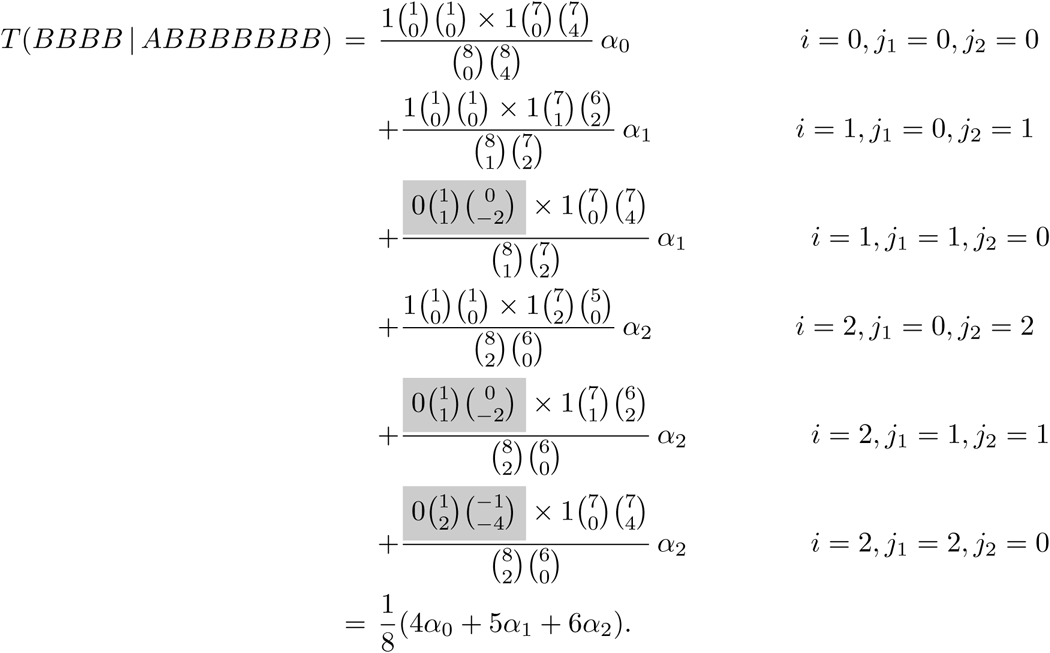

Note that *δ*_*jk*_ = 0 whenever *j*_*k*_ < max(0, *m*_*k*_ - *n*_*k*_) or *j*_*k*_ > min(*n*_*k*_, *m*_*k*_/2). The coefficients of the above terms with a gray background are equal to zero.

### B Derivation of *α*_*i*_ **under octosomic inheritance**

In this example, we show how to derive the values of alpha for octosomic inheritance in both CES and PES.

For the case of CES, we firstly calculate the values of *P*_*j*_ for each possible *j*. Because there are *v*/2 chromosomes, the maximum value of *j* is ⌊*v*/4⌋. Note that in this case, we have *v* = 8, then ⌊*v*/4⌋ = 2 and *j* = 0, 1, 2. Now, by Equation (4), we obtain

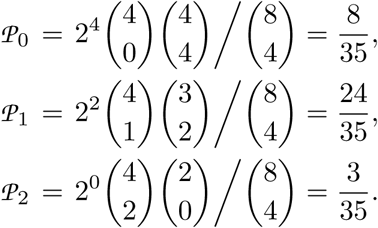

Secondly, we calculate the value of *α*_*i*_ under CES where 0 ⩽ *i* ⩽ ⌊*v*/4⌋, i.e. 0 ⩽ *i* ⩽ 2. By Equation (5), it follows

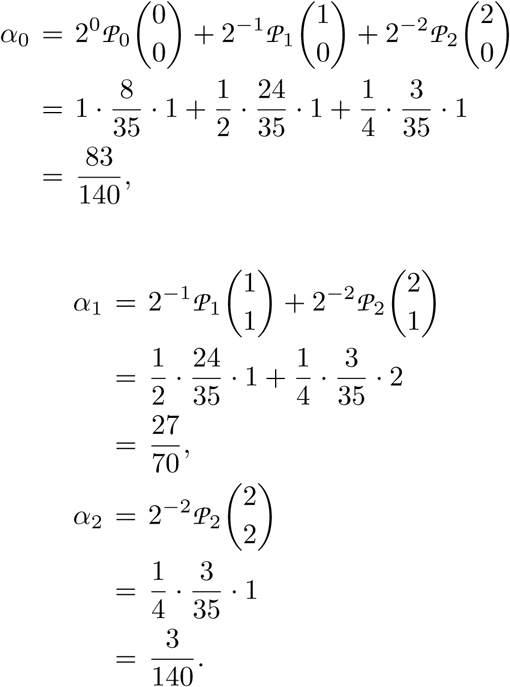

For the case of PES, we still have *v* = 8, then ⌊*v*/4⌋ = 2 and the values of *P*_0_, *P*_1_ and *P*_2_ in Equation (6) are just those values in the case of CES. Now, according to Equation (6), we have

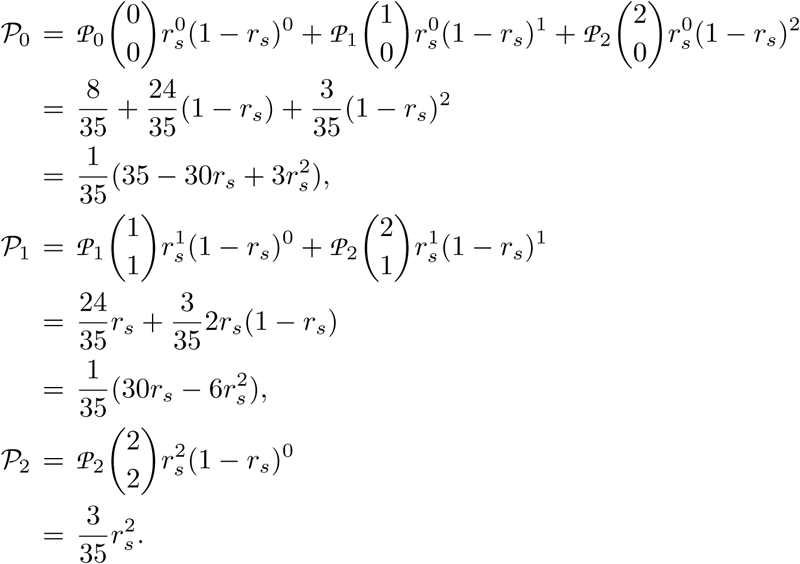

Further, because of Equation (5), we derive the value of *α*_*k*_ (*k* = 0, 1, 2) under PES as follows:

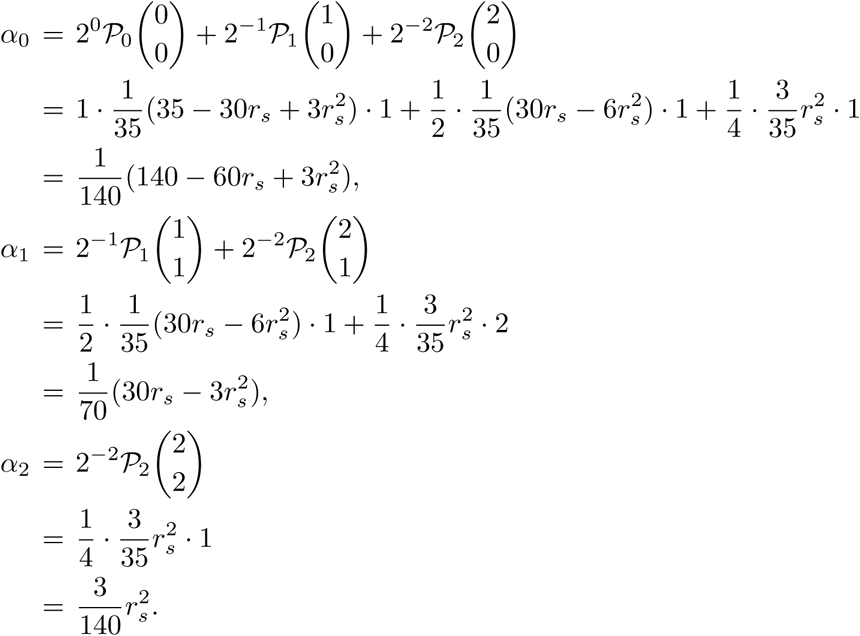

### C Derivation of GFG and GFZ with nonlinear method

Here, we use the tetrasomic inheritance at a triallelic locus under equilibrium to derive the GFG by using a non-linear method as an example. Under these conditions, we have *v* = 4 and *h* = 3. Because 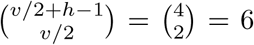 and 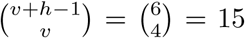, there are 6 gamete genotypes and 15 zygote genotypes, and so Equations (8) and (9) determine 6 and 15 equations, respectively.

(i) Simulating meiosis. We denote *AA* and *AB* for two gamete genotypes, and *P*_*AA*_ and *P*_*AB*_ for their frequencies, and so on. Similarly, denote *AABB* and *ABBC* for two zygote genotypes, and *P*_*AABB*_ and *P*_*ABBC*_ for their frequencies, and so on. Now, by Equation (8), the GFG can be established as follows:

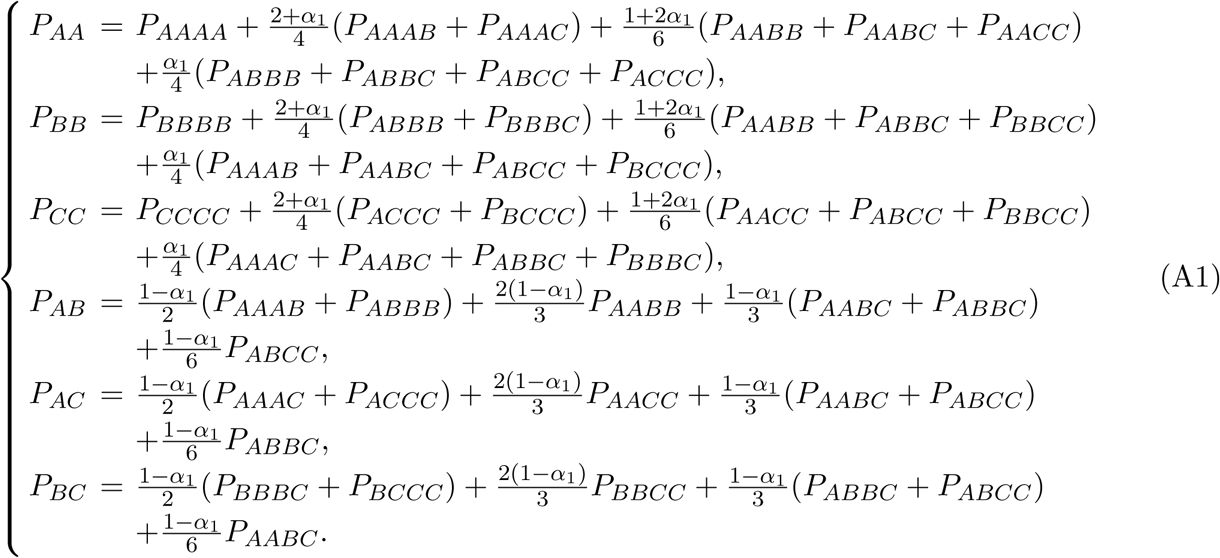

(ii) Simulating fertilization. By Equation (9), the GFZ can be established as follows:

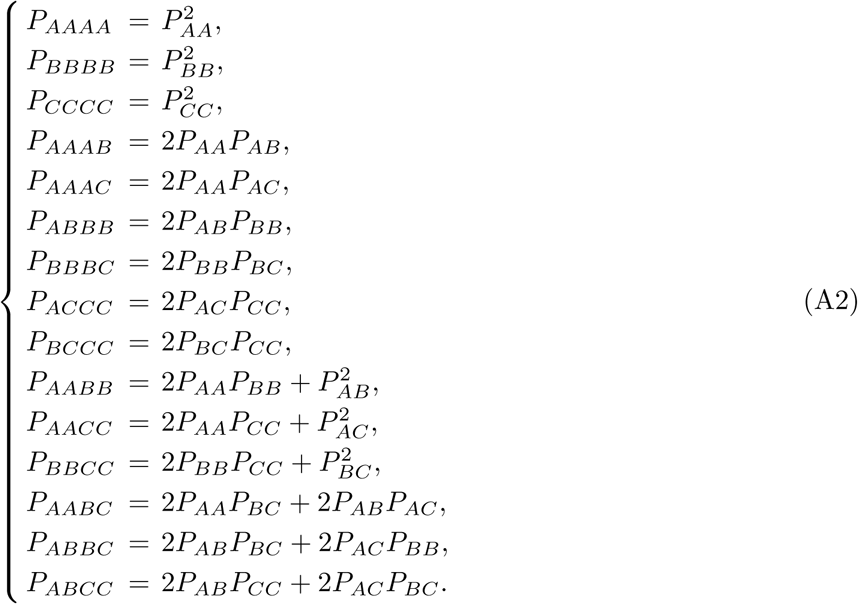

Now, substituting Equation (A2) into Equation (A1), the GFZ are eliminated, and a system of nonlinear equations with 6 equations and 6 unknown is obtained (whose expressions are more complex and omitted). On the other hand, the process that transform the allele frequencies into GFG can described by the linear substitution

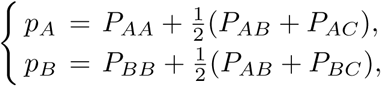

where *p*_*A*_, *p*_*B*_ and *p*_*C*_ are the allele frequencies with *p*_*A*_ + *p*_*B*_ + *p*_*C*_ = 1. Combining the linear substitution with the system of non-linear equations mentioned above, we still obtain a system of non-linear equations with 8 equations, 6 unknowns (i.e. *P*_*AA*_, *P*_*AB*_, *P*_*AC*_, *P*_*BB*_, *P*_*BC*_ and *P*_*CC*_) and 2 parametric variables (i.e. *p*_*A*_ and *p*_*B*_). The solution that is with *p*_*A*_ and *p*_*B*_ as the parametric variables is unique and is shown as follows:

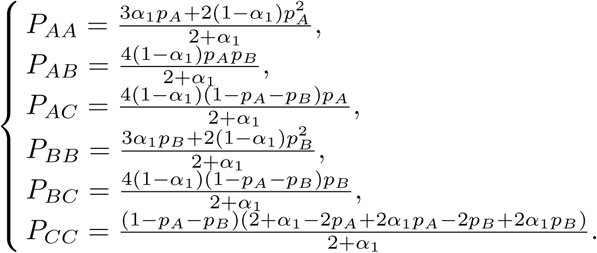

Therefore the generalized form for GFG at equilibrium can be directly written as follows:

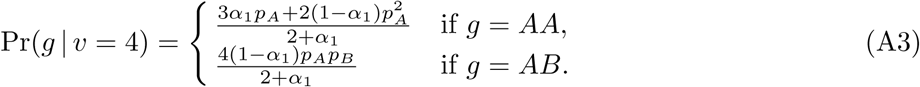

Using Equation (9), we can derive the generalized form for GFZ at equilibrium as follows:

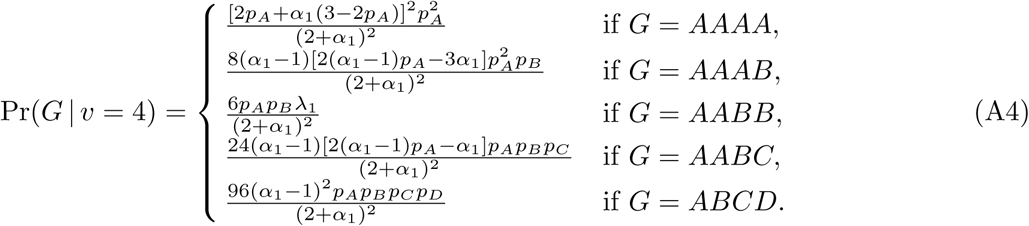

Where 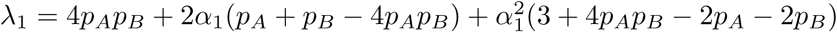.

### D GFG and GFZ in hexasomic inheritance

The generalized form of GFG for hexasomic inheritance at equilibrium derived from the linear method is given by

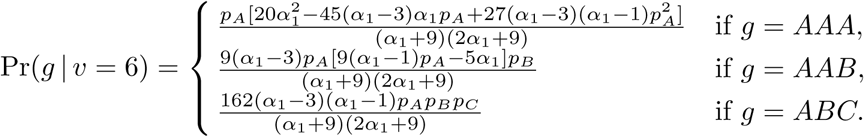

Where *AAA, AAB* and *ABC* are the genotypic patterns of *g*. With Equation (9), the generalized form of GFZ for hexasomic inheritance at equilibrium is given by

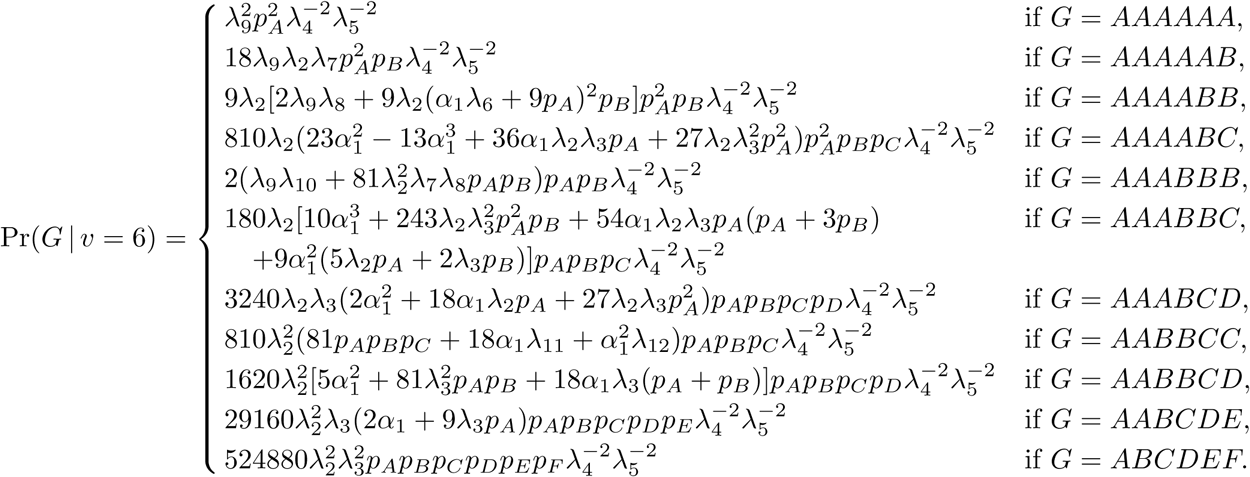

Where 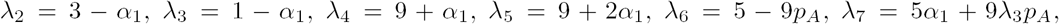 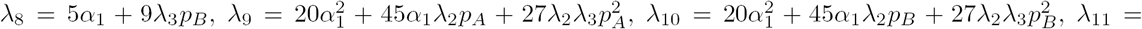 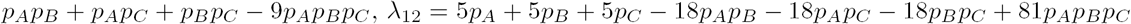.

### E Derivation of co-ancestry coefficients in different relationships

The expression of co-ancestry coefficient *θ* in mating individuals can be obtained by taken the weighted average of co-ancestry coefficient between different relationships, where the weight is the frequencies of those relationships in mating individuals. We first derive the co-ancestry coefficients in the following four relationships.

In selfing, the individual self-fertilizes. For a pair of alleles sampled from an individual with replacement, the probability is 1/*v* if the same allele is sampled twice and these are indeed IBD; otherwise the probability is *F*. Hence 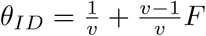.

In backcrossing, the offspring is fertilized by, or fertilizes, its parent (says *f*). Let *m* be the other parent. Denote *g*_*f*_ and *g*_*m*_ for the two gametes which are respectively produced by *f* and *m* to form an offspring. The allele pairs between the offspring and *f* can be classified into two categories: (i) between *g*_*f*_ and *f*. In this case, for each allele pair, the probability that the two alleles are IBD is equal to *θ*_*ID*_; (ii) between *g*_*m*_ and *f*. In this case, for each allele pair, the probability that the two alleles are IBD alleles is equal to *θ*, i.e. the co-ancestry coefficient between mating individuals. Hence *θ*_*PO*_ = (*θ*_*ID*_ + *θ*)/2.

In matings between full-siblings (says *a* and *b*), it is assumed that the parents are *f* and *m*, and let *g*_*af*_ and gam be the gametes forming *a*, where *g*_*af*_ is produced by *f* and *g*_*am*_ is produced by *m*. Similarly *g*_*bf*_ and *g*_*bm*_ denote the gametes forming *b*. For each pair of alleles, the probability that the two alleles are IBD between *g*_*af*_-*g*_*bf*_ is *θ*_*ID*_, the same as that between *g*_*am*_-*g*_*bm*_; and the probability that they are IBD between *g*_*af*_ -*g*_*bm*_ is *θ*, the same as that between *g*_*am*_-*g*_*bf*_. Hence *θ*_*FS*_ = (*θ*_*ID*_ + *θ*)/2.

In matings between nonrelatives, *θ*_*UN*_ = 0.

Second, we derive the *F* in population with a selfing ratio *s* as an example. Assuming a proportion *s* of individuals is produced by selfing and the remaining proportion 1 − *s* of individuals is produced from matings between nonrelatives, then *θ* = *s θ*_*ID*_ + (1 − *s*)*θ*_*UN*_. Because *θ*_*UN*_ = 0, this expression can be simplified into *θ* = *s θ*_*ID*_. By substituting this expression into Equation (11), we obtain inbreeding coefficient *F* at equilibrium: 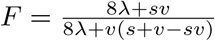.

